# Citizen-science surveillance of triazole-resistant *Aspergillus fumigatus* in UK residential garden soils

**DOI:** 10.1101/2021.07.08.451577

**Authors:** Jennifer M. G. Shelton, Roseanna Collins, Christopher B. Uzzell, Asmaa Alghamdi, Paul S. Dyer, Andrew C. Singer, Matthew C. Fisher

## Abstract

Compost is an ecological niche for *Aspergillus fumigatus* due to its role as a decomposer of organic matter and its ability to survive the high temperatures associated with the composting process. Subsequently, composting facilities are associated with high levels of *A. fumigatus* spores that are aerosolised from compost and cause respiratory illness in workers. In the UK, gardening is an activity enjoyed by individuals of all ages and it is likely that they are being exposed to *A. fumigatus* spores when handling commercial compost or compost they have produced themselves. In this study, 246 citizen scientists collected 509 soil samples from locations in their garden in the UK, from which were cultured 5,174 *A. fumigatus* isolates. Of these isolates, 736 (14%) were resistant to tebuconazole: the third most-sprayed triazole fungicide in the UK, which confers cross-resistance to the medical triazoles used to treat *A. fumigatus* lung infections in humans. These isolates were found to contain the common resistance mechanisms in the *A. fumigatus cyp51A* gene TR_34_/L98H or TR_46_/Y121F/T289A, and less common resistance mechanisms TR_34_, TR_53_, TR_46_/Y121F/T289A/S363P/I364V/G448S and (TR_46_)^2^/Y121F/M172I/T289A/G448S. Regression analyses found that soil samples containing compost were significantly more likely to grow susceptible and tebuconazole-resistant *A. fumigatus* than those that did not, and that compost samples grew significantly higher numbers of *A. fumigatus* than other samples.

**Importance:** These findings highlight compost as a potential health hazard to individuals with pre-disposing factors to *A. fumigatus* lung infections, and a potential health hazard to immunocompetent individuals who could be exposed to sufficiently high numbers of spores to develop infection. This raises the question of whether compost bags should carry additional health warnings regarding inhalation of *A. fumigatus* spores, whether individuals should be advised to wear facemasks whilst handling compost or whether commercial producers should be responsible for sterilising compost before shipping. The findings support increasing public awareness of the hazard posed by compost and investigating measures that can be taken to reduce the exposure risk.

## Introduction

The fungus *Aspergillus fumigatus* plays an important role in the environment as a decomposer, recycling nutrients from decaying plant matter into the soil. This highly sporulating mould is commonly found in woodchip piles, compost from household waste, sewage, sludge and mouldy hay^1^, where its thermotolerance enables it to proliferate during the thermogenic phase of composting when temperatures reach 40-60°C^2^. The small size of *A. fumigatus* spores (2-3 μm) and their hydrophobicity means they are easily aerosolised and transported on air currents, making *A. fumigatus* a globally ubiquitous fungus^3^. Exposure to this mould is medically important and it is estimated that humans inhale several hundred *A. fumigatus* spores per day^4^, which can trigger an immunoinflammatory response resulting in severe asthma with fungal sensitisation (SAFS) or allergic bronchopulmonary aspergillosis (ABPA)^5^. The size of the spores allows them to bypass mucociliary clearance in the lung^6^ and their thermotolerance enables them to establish in lung cavities if not cleared first by the innate immune system, resulting in chronic pulmonary aspergillosis (CPA). Individuals who are immunocompromised - due to treatment with immunosuppressants, chemotherapy, diabetes, rheumatoid arthritis or HIV/AIDS infection - or who have impaired lung function from chronic obstructive pulmonary disease (COPD), tuberculosis (TB), severe influenza or COVID-19 infection or cystic fibrosis (CF) are at greatest risk of CPA. Invasive aspergillosis (IA) occurs if the immune system is unable to prevent spores from entering the bloodstream, which is a life-threatening infection associated with ~58% survival^7^. It was estimated that in the UK in 2011 there were ~178,000 individuals living with ABPA, 3,600 with CPA and 2,900 with IA, plus an additional 377-1,345 cases of IA in critical care patients^8^. The number of patients in the UK presenting with infections that are resistant to one or more of itraconazole (ICZ), voriconazole (VCZ) and posaconazole (PCZ) – the frontline triazole drugs for treating aspergillosis – has risen from 3-7% between 1999 and 2001 to 14-20% between 2007 and 2009^9^. Triazole-resistant infections are associated with treatment failure, salvage therapy with more toxic antifungals and increased case fatality rates (CFR), with CFRs up to 88% reported for triazole-resistant IA^10^.

Triazole-resistance is most commonly caused by polymorphisms in the *cyp51A* gene, which results in increased production of, or configurational changes in, lanosterol-14a-demethylase; an enzyme involved in ergosterol biosynthesis and the binding target of triazole drugs. An environmental route for the acquisition of triazole-resistant infections has been proposed due to the increase of infections caused by *A. fumigatus* isolates with a tandem repeat (TR) in the promoter region of *cyp51A* coupled with single nucleotide polymorphisms (SNPs) in the coding region leading to amino acid substitutions in the protein, which are frequently recovered from air and soil samples globally^11^. This is likely due to the use of fungicides epoxiconazole, tebuconazole, propiconazole, difenoconazole and bromuconazole, which have similar molecular structures to the medical triazoles and show cross-resistance^12^. In 2008, these were the second, third, sixth, ninth and seventeenth most sprayed triazoles in agriculture in the UK, respectively^13^. In agriculture, triazoles are applied to wheat, beans, carrots, oilseed rape, soft fruits and vines; in horticulture, they are used to sterilise bulbs and to control fungal diseases in lawns and ornamentals; and in industry they are used as wood preservatives and antifouling agents in leather, paper, textiles, paints and adhesives^13^.

The UK government is committed to reducing carbon dioxide emissions by diverting waste from landfill and incineration to composting^14^, and compost features in the government’s Food 2030 strategy for improving the productive capacity of soil^15^. Compost producers accept input material from agriculture, horticulture, forestry, wood and paper processing, leather and textiles industries, household and garden waste, which are highly likely to contain triazole residues. In 2007, 90% of composting facilities in the UK produced compost in open windrows^16^; where organic waste is shredded, mixed and placed in uncovered rows that are turned regularly during the composting process to improve oxygenation of the waste and to distribute heat and moisture. Composting facilities are known to produce large numbers of *A. fumigatus* spores^16–23^, with resulting negative health impacts on compost handlers^24–32^, and there is evidence from the Netherlands that composting material also produces large numbers of triazole-resistant spores^33,34^. In 2017, UK households spent approximately £450 million on compost^35^ and apply it more liberally to their gardens at 300 tonnes per hectare (t/ha) than the 50 t/ha applied to agricultural land^36^. Furthermore, more than a third of households with access to a garden report composting their garden and/or kitchen waste^37^. This means that a substantial proportion of the UK population is handling compost on a regular basis, with potential exposure to high levels of *A. fumigatus* spores that may have developed triazole-resistance from composts that contain triazole residues. Indeed, there have been reports of hypersensitivity pneumonitis^38^ and IA^39–43^ in apparently immunocompetent individuals following gardening activities, however, no clinical links following exposure to triazole resistant spores have been documented.

The aims of this study were to a) determine the numbers of triazole-susceptible and resistant *A. fumigatus* spores in soil samples collected from residential gardens in the UK, b) characterise the *cyp51A* polymorphisms responsible for resistance, and c) find environmental variables associated with presence/numbers of *A. fumigatus* spores in soil samples. In order to simultaneously sample a wide range of UK gardens, we were assisted by a network of citizen-scientists trained in the collection of samples that may contain *A. fumigatus*. Our aim was to ascertain whether gardening activities may lead to exposure to triazole-resistant genotypes of this mould that could present a risk to susceptible individuals. Based on our findings, we present thoughts on how these exposure risks in susceptible individuals might be mitigated.

## Methods

### *Culturing* Aspergillus fumigatus *from residential garden soil samples*

The soil samples from which *A fumigatus* isolates were cultured for this study were collected as part of a citizen science project undertaken in June 2019, which involved 246 volunteers in the UK collected a total of 509 soil samples from different locations in their gardens^44^. Participants indicated on a questionnaire whether samples were collected from a border, pot or planter, compost heap, bag of manure or bag of compost. Upon receipt, 2 g of each soil sample was suspended in 8 ml of buffer (0.85% NaCl and 0.01% Tween 20 in distilled water), shaken vigorously and left to settle for 30 minutes. One aliquot of 200 μl from the surface of the buffer was spread onto a plate containing sabouraud dextrose agar (SDA), penicillin (200 mg/L) and streptomycin (400 mg/L) and a second aliquot of 200 μl was spread onto a plate containing SDA, penicillin (200 mg/L), streptomycin (400 mg/L) and tebuconazole (6 mg/L). Both plates were incubated at 37°C for 48 hours, the number of *A. fumigatus* colonies on each plate recorded, and the colonies growing on the plate containing tebuconazole were picked into tubes containing mould preservation solution (0.2% agar and 0.05% Tween 20 in dionized water) and stored at 4°C. These isolates were subsequently cryopreserved in 50% glycerol solution and were DNA extracted as detailed in Boyle *et al.* (2004)^45^.

### *Sequencing of* A. fumigatus cyp51A *gene*

The promoter region of *cyp51A* was amplified using forward primer 5’-GGACTGGCTGATCAAACTATGC-3’ and reverse primer 5’-GTTCTGTTCGGTTCCAAAGCC-3’ and the following PCR conditions: 95°C for five minutes; 30 cycles of 98°C for 20 seconds, 65°C for 30 seconds and 72°C for 30 seconds; followed by 72°C for five minutes. Amplicons were visualised by gel electrophoresis and samples with visible bands were sent for sequencing using the forward primer. The coding region of *cyp51A* was amplified using forward primer 5’-ATGGTGCCGATGCTATGG-3’ and reverse primer 5’-CTGTCTCACTTGGATGTG-3’ and the following PCR conditions: 94°C for two minutes; 35 cycles of 94°C for 30 seconds, 60°C for 45 seconds and 72°C for 45 seconds; followed by 72°C for five minutes. Amplicons were visualised by gel electrophoresis and samples with visible bands were sent for sequencing using the Sanger chain termination method in two segments using the primers 5’-TACGTTGACATCATCAATCAG-3’ and 5’-GATTCACCGAACTTTCAAGGCTCG-3’. Sequences were aligned using Molecular Evolutionary Genetics Analysis (MEGA) software (Penn State University, US).

### Identification of isolates

For isolates that failed to sequence using the primers for the promoter and coding regions of *cyp51A*, part of the beta-tubulin gene was sequenced using forward primer 5’-AATTGGTGCCGCTTTCTGG-3’ and reverse primer 5’-AGTTGTCGGGACGGAATAG-3’ and the following PCR conditions: 94°C for 3 minutes; 30 cycles of 94°C for 15 seconds, 55°C for 30 seconds, 68°C for 30 seconds; followed by 68°C for 3 minutes. Amplicons were visualised by gel electrophoresis and samples with visible bands were sent for sequencing using the forward primer. Basic Local Alignment Search Tool (BLAST) was used to align the sequences to those in the National Center for Biotechnology Information (NCBI; Bethesda, US) to identify the isolate.

### *Environmental variables that may influence growth of* Aspergillus fumigatus

Table 1 details the environmental variables that were ascertained for the locations in the UK from which soil samples were collected, on the date when sampling occurred, and the source from which the data were obtained.

**Table 1:**
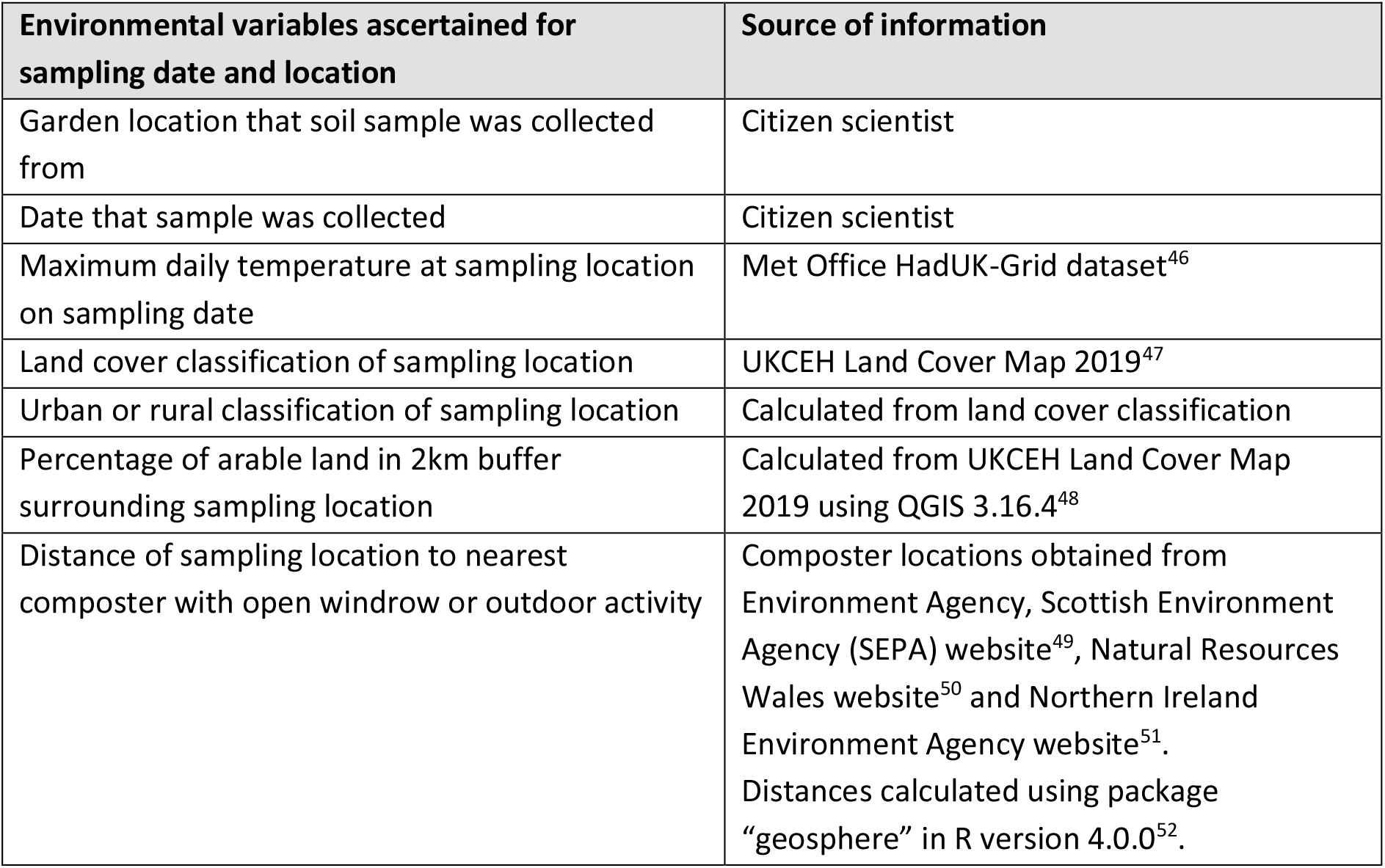
Environmental variables obtained for soil sampling locations and dates and the sources they were obtained from.

### Generalised linear models

Generalised linear models (GLMs) were run using R version 4.0.0 to find associations between the environmental variables in Table 1 and 1) the likelihoods of a sample growing susceptible or triazole-resistant *A. fumigatus*, and 2) the number of susceptible or triazole-resistant *A. fumigatus* colonies grown from a sample. Growth of susceptible or triazole-resistant *A. fumigatus* from a sample was categorised as 0/1 and logistic regressions (“glm” function; family = “binomial”) were performed. The numbers of susceptible and triazole-resistant *A. fumigatus* colonies grown from samples were over-dispersed; therefore negative binomial regressions (library “MASS”; “glm.nb” function) were performed. Environmental variables were included in the regression model based on a significant improvement on the null model, as determined by analysis of variance (ANOVA) using chi-squared test. Results were considered significant when *p* <= 0.05. The regression model with the best fit was chosen based on a reduced Akaike information criterion (AIC) score and a significant improvement on the null model.

## Results

### *Susceptible and tebuconazole-resistant* A. fumigatus *in soil samples*

Of the 509 soil samples collected, 327 (64%) samples between them grew 5,174 *A. fumigatus* isolates and 101 (20%) samples grew 736 tebuconazole-resistant isolates (Table 2). The majority of samples (*n* = 451; 89%) were assigned a single location in the garden from which they were collected, whereas the remainder were assigned multiple locations. These multiple locations occurred when a border or pot/planter had recently been topped up with manure or compost. The concentration of spores averaged across the samples that grew *A. fumigatus* was 316 CFU/g, which ranged from 0 CFU/g in the sample collected from a border plus manure bag to 600 CFU/g in the sample collected from a manure bag. The concentration of spores averaged across the samples that grew tebuconazole-resistant *A. fumigatus* was 146 CFU/g, which ranged from 0 CFU/g in samples collected from several garden locations to 214 CFU/g in samples collected from compost heaps. Figure 1 shows the geographical locations in the UK that soil samples were collected from.

**Table 2:** A breakdown of the number of soil samples collected, the number and percentage of soil samples that grew susceptible and tebuconazole-resistant *Aspergillus fumigatus*, the numbers of susceptible and tebuconazole-resistant *A. fumigatus* isolates grown and the average colony forming unit per gram (CFU/g) across samples that grew susceptible and tebuconazole-resistant *A. fumigatus* by the location(s) in the garden that the soil sample was collected from.

| Location in garden that soil sample was collected from | Number of soil samples | Samples that grew *A. fumigatus* (% of samples) | Samples that grew tebuconazole-resistant *A. fumigatus* (% of samples) | Number of *A. fumigatus* isolates grown | Average *A. fumigatus* CFU/g | Number of tebuconazole-resistant *A. fumigatus* isolates grown (% of *A. fumigatus* isolates) | Average tebuconazole-resistant *A. fumigatus* CFU/g |
| --- | --- | --- | --- | --- | --- | --- | --- |
| Border | 206 | 99 (48) | 19 (9) | 1,009 | 204 | 121 (12) | 127 |
| + compost bag | 7 | 4 (57) | 1 (14) | 56 | 280 | 8 (14) | 160 |
| + compost heap | 5 | 3 (60) | 0 (0) | 44 | 293 | 0 (0) | 0 |
| + manure bag | 1 | 0 (0) | 0 (0) | 0 | 0 | 0 (0) | 0 |
| + manure bag + compost heap | 1 | 1 (100) | 0 (0) | 1 | 20 | 0 (0) | 0 |
| Compost bag | 49 | 44 (90) | 20 (41) | 993 | 451 | 137 (14) | 137 |
| Compost heap | 80 | 58 (73) | 27 (34) | 1,464 | 505 | 289 (20) | 214 |
| Manure bag | 1 | 1 (100) | 0 (0) | 30 | 600 | 0 (0) | 0 |
| Pot/planter | 115 | 79 (69) | 26 (23) | 1,005 | 254 | 130 (13) | 98 |
| + compost bag | 38 | 33 (87) | 8 (21) | 529 | 321 | 51 (15) | 128 |
| + compost bag + compost heap | 3 | 2 (67) | 0 (0) | 21 | 210 | 0 (0) | 0 |
| + compost bag + manure bag | 2 | 2 (100) | 0 (0) | 4 | 40 | 0 (0) | 0 |
| + compost heap | 1 | 1 (100) | 0 (0) | 18 | 360 | 0 (0) | 0 |
| **Total** | **509** | **327 (64)** | **101 (20)** | **5,174** | **316** | **736 (14)** | **145** |

**Figure 1:**
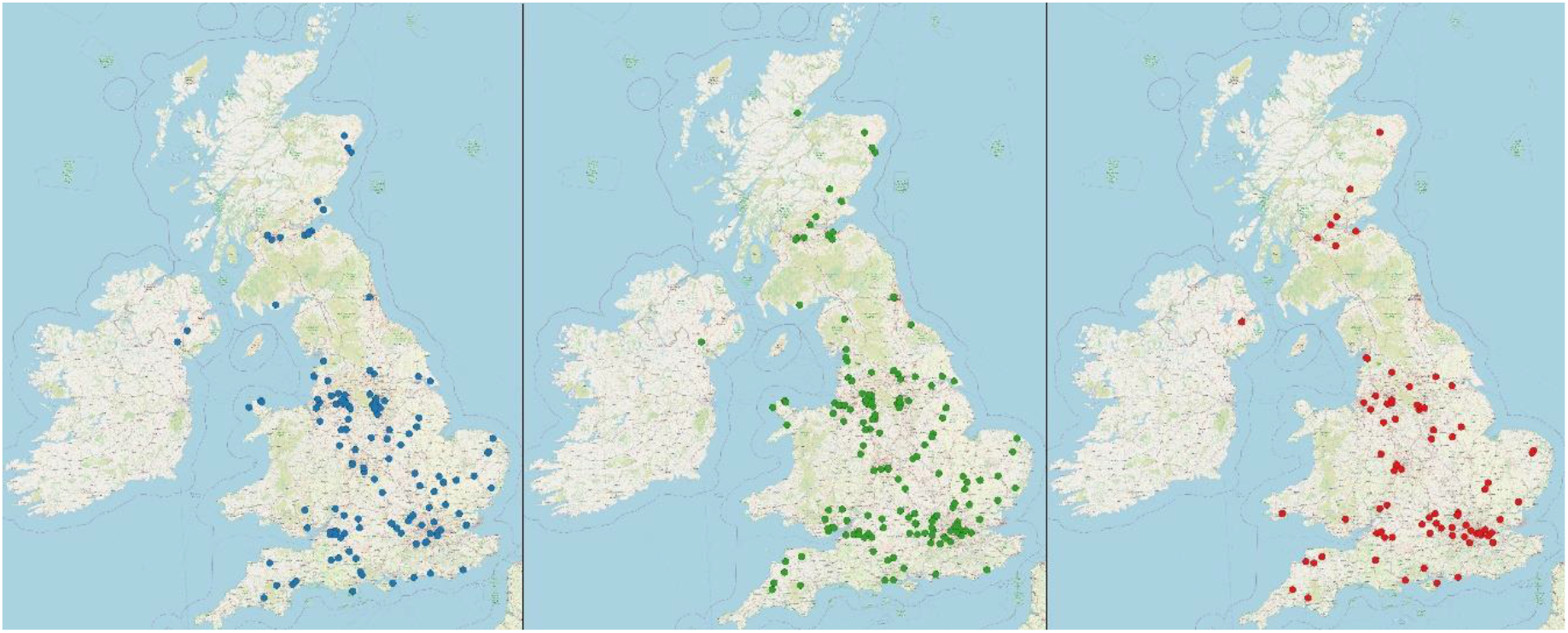
Geographical locations in the UK that soil samples were collected from by citizen scientists. Blue dots indicate samples that did not grow *Aspergillus fumigatus*, green dots indicate samples that grew *Aspergillus fumigatus* and red dots indicate samples that grew tebuconazole-resistant *A. fumigatus*. Base maps created using data obtained from OpenStreetMap (CC BY-SA 4.0); URL: https://www.openstreetmap.org.

### Cyp51A *polymorphisms in tebuconazole-resistant* A. fumigatus *isolates*

Of the 736 tebuconazole-resistant *A. fumigatus* isolates, 93 (13%) failed to re-grow from refrigerated storage for cryopreservation and DNA extraction. In the 643 isolates that re-grew, TR34/L98H was detected in 542 (85%), TR_46_/Y121F/T289A in 16 (3%), TR_53_ in two, (TR_48_)^2^/Y121F/M172I/T289A/G448S in one and no *cyp51A* polymorphisms were detected in 27 (4%) isolates. 14 isolates failed to sequence with the *cyp51A* promoter and coding region primers and beta-tubulin sequencing confirmed their identities as *A. fischeri* (*n* = 8), *A. fumigatus* (*n* = 2), *A. oerlinghausensis* (*n* = 3) and unknown (*n* = 1). Uncommon polymorphisms detected were TR_34_ without accompanying amino acid substitutions in three isolates, (TR_34_)^2^/L98H in one isolate and (TR_130_)^3^/D430G in four isolates. The remaining isolates contained one or more amino acid substitutions in cyp51A, with or without accompanying TRs (Table 3). Further details of the tebuconazole-resistant *A. fumigatus* isolates can be found in Supplementary Table 1.

**Table 3:** *cyp51a* polymorphisms for the 636 tebuconazole-resistant *Aspergillus fumigatus* isolates, by garden location they were collected from. ^a^ Samples that failed to amplify with the *cyp51A* promoter and coding region primers were sequenced using beta-tubulin primers for fungal identification. B=border, CB=compost bag, CH=compost heap, MB=manure bag, PP=pot/planter.

| Tandem repeat in *cyp51A* promoter region | Amino acid substitution(s) in *cyp51A* | Location in garden that soil sample was collected from |  |  |  |  |  | Total: | Medical triazole susceptibility |
| --- | --- | --- | --- | --- | --- | --- | --- | --- | --- |
|  |  | B | B+CB | CB | CH | PP | PP+CB |  |  |
| - | - | 3 |  |  | 1 | 23 |  | 27 |  |
| - | C270R |  |  |  |  | 1 |  | 1 |  |
| - | I242V |  |  |  |  | 5 |  | 5 | 53 |
| TR<sub>34</sub> | - |  |  |  | 1 | 1 | 1 | 3 | 54 |
| TR<sub>34</sub> | L98H | 82 | 7 | 117 | 237 | 65 | 34 | 542 | 55 |
| TR<sub>34</sub> | L98H/Q191E |  |  | 1 |  |  |  | 1 |  |
| TR<sub>34</sub> | L98H/R196L |  | 1 |  |  |  |  | 1 |  |
| TR<sub>34</sub> | L98H/K240R | 1 |  |  |  | 1 |  | 2 |  |
| TR<sub>34</sub> | L98H/T289A/I364V/G448S |  |  |  | 6 |  |  | 6 | 56 |
| TR<sub>34</sub> | L98H/K372R |  |  |  | 1 |  |  | 1 |  |
| TR<sub>34</sub> | L98H/P394R |  |  | 1 |  |  |  | 1 |  |
| TR<sub>34</sub> | L98H/F404C/F459S/A460S |  |  | 1 |  |  |  | 1 |  |
| TR<sub>34</sub> | L98H/F404V |  |  | 1 |  |  |  | 1 |  |
| TR<sub>34</sub> | L98H/N406D |  |  | 1 |  |  |  | 1 |  |
| TR<sub>34</sub> | L98H/N406M |  |  |  | 1 |  |  | 1 |  |
| TR<sub>34</sub> | L98H/K421R |  |  | 1 |  |  |  | 1 |  |
| TR<sub>34</sub> | L98H/P443L |  |  |  |  | 2 |  | 2 |  |
| TR<sub>34</sub> | L98H/A460S |  |  |  |  | 1 |  | 1 |  |
| TR<sub>34</sub> | L98H/D481N |  |  | 1 |  |  | 1 | 2 |  |
| (TR<sub>34</sub>)<sup>2</sup> | L98H | 1 |  |  |  |  |  | 1 |  |
| TR<sub>46</sub> | Y121F/M178W/T289A/S363P/I364V/G448S |  |  |  | 1 |  |  | 1 |  |
| TR<sub>46</sub> | Y121F/T289A |  |  |  | 15 |  | 1 | 16 |  |
| TR<sub>46</sub> | Y121F/T289A/S363P/I364V/G448S |  |  |  | 4 |  |  | 4 | 57 |
| (TR<sub>46</sub>)<sup>2</sup> | Y121F/M172I/T289A/G448S |  |  |  | 1 |  |  | 1 | 34 |
| TR<sub>53</sub> | - | 1 |  |  | 1 |  |  | 2 | 58 |
| (TR<sub>130</sub>)<sup>3</sup> | D430G |  |  |  | 4 |  |  | 4 | 59 |
| Failed to sequence | Failed to sequence<sup>a</sup> | 3 |  |  | 3 | 8 |  | 14 |  |
| Total: |  | 91 | 8 | 124 | 276 | 107 | 37 | 643 |  |

### *Environmental variables influencing growth and numbers of* A. fumigatus *colonies*

#### *Growth of* A. fumigatus *from soil samples*

Eight samples were excluded from the logistic regression with growth of *A. fumigatus* as the outcome because the SDA plates were too contaminated to determine presence of *A. fumigatus*, which left 501 samples in the analysis. Location in the garden from which the soil sample was collected was the only variable that significantly affected whether a sample grew *A. fumigatus* (χ^2^ = 67.3, df = 12, *p* < 0.01). The odds ratios and p-values from the logistic regression model are shown in Table 4. Samples collected from a compost bag, compost heap, pot/planter and pot/planter plus compost bag had significantly increased odds of growing *A. fumigatus* (*p* < 0.01) compared to samples collected from a border. There were no significant changes in odds of growing *A. fumigatus* from other sampling locations.

**Table 4:** Odds ratios, confidence intervals and p-values from logistic regression model using location in garden that sample was collected from as an explanatory variable for whether samples (*n* = 501) grew *Aspergillus fumigatus*. Significant results (p <= 0.05) are highlighted in bold. ^a^ Insufficient data to calculate odds ratio and confidence intervals.

| Environmental variable | Odds ratio (95% CI) | Pr(>\|z\|) |
| --- | --- | --- |
| Location in garden sampled from: |  |  |
| Border (baseline) |  |  |
| + compost bag | 1.43 (0.31-7.40) | 0.64 |
| + compost heap | 1.61 (0.26-10.24) | 0.61 |
| + manure bag | -<sup>a</sup> | 0.99 |
| + manure bag + compost heap | -<sup>a</sup> | 0.99 |
| Compost bag | **15.70 (5.50-66.19)** | **<0.01** |
| Compost heap | **3.45 (1.93-6.40)** | **<0.01** |
| Manure bag | -<sup>a</sup> | 0.99 |
| Pot/planter | **2.42 (1.50-3.95)** | **<0.01** |
| + compost bag | **7.07 (2.88-21.28)** | **<0.01** |
| + compost bag + compost heap | 2.14 (0.20-46.50) | 0.53 |
| + compost bag + manure bag | -<sup>a</sup> | 0.98 |
| + compost heap | -<sup>a</sup> | 0.99 |

### *Number of* A. fumigatus *colonies* grown *from soil samples*

The first negative binomial regression was run on the 335 samples that grew *A. fumigatus*. The only variable found to significantly affect the number of *A. fumigatus* colonies grown from a sample was garden location from which the sample was collected (χ^2^ = 50.8, df = 11, *p* < 0.01). In the regression model, samples collected from compost bag (*p* < 0.01), compost heap (*p* < 0.01) and pot/planter plus compost bag (*p* = 0.02) grew significantly more *A. fumigatus* colonies than samples collected from borders. Samples collected from a pot/planter plus compost bag plus manure bag grew fewer *A. fumigatus* colonies than samples collected from borders, although this reduction was marginally significant (*p* = 0.05).

### *Growth of tebuconazole-resistant* A. fumigatus *from soil samples*

All 509 soil samples were included in the logistic regression with growth of tebuconazole-resistant *A. fumigatus* as the outcome. The only variable found to significantly affect whether a sample grew tebuconazole-resistant *A. fumigatus* was garden location from which the sample was collected (χ^2^ = 43.0, df = 12, *p* < 0.01). The odds ratios and p-values from the logistic regression model are shown in Table 5. Samples collected from a compost bag, compost heap, pot/planter and pot/planter plus compost bag had significantly increased odds of growing tebuconazole-resistant *A. fumigatus* (*p* < 0.01) compared to samples collected from a border. There were no significant changes in odds of growing tebuconazole-resistant *A. fumigatus* from other sampling locations.

**Table 5:**
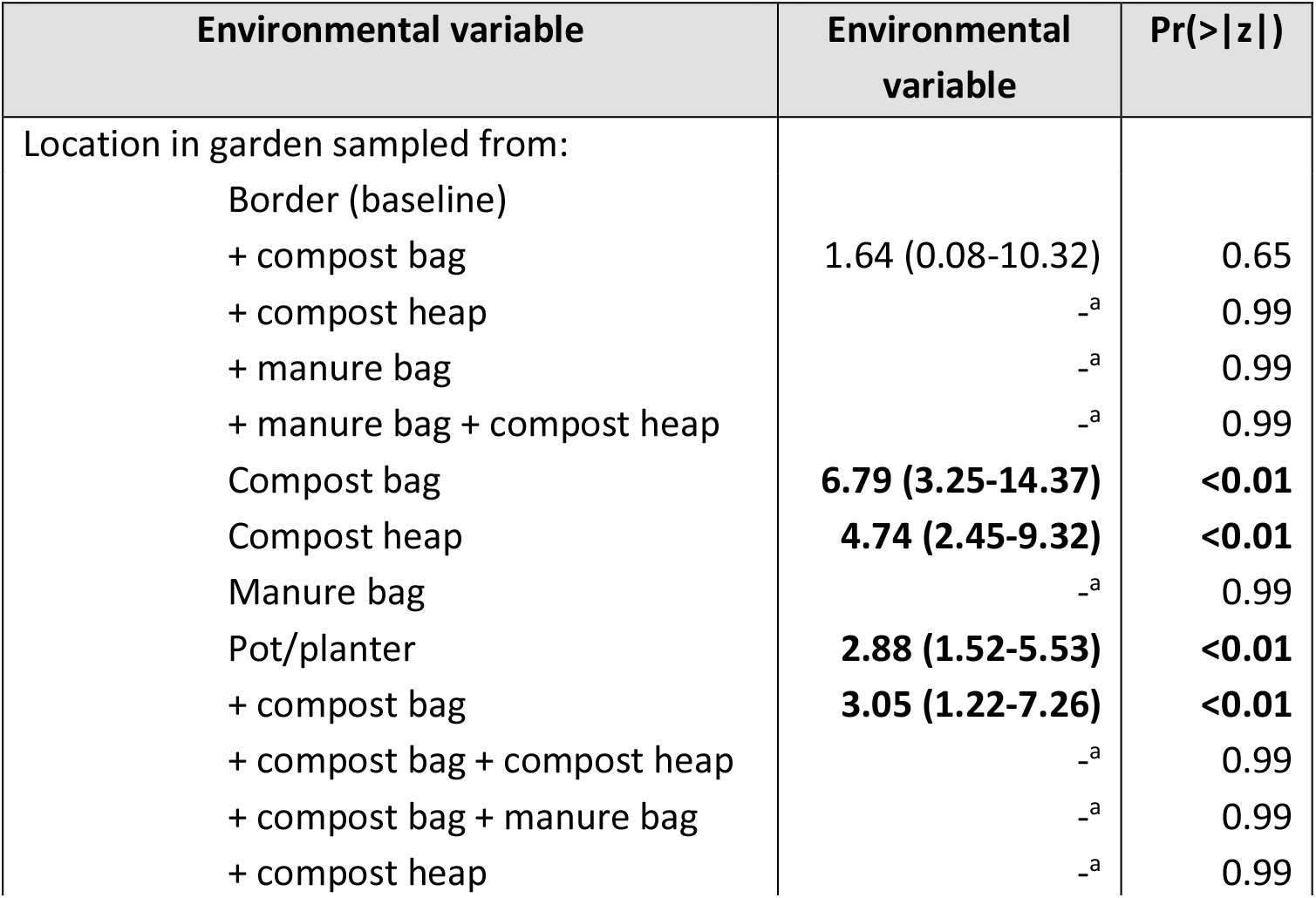
Odds ratios, confidence intervals and p-values from logistic regression model using location in garden that sample was collected from as an explanatory variable for whether samples (*n* = 509) grew tebuconazole-resistant *Aspergillus fumigatus*. Significant results (p <= 0.05) are highlighted in bold. ^a^ Insufficient data to calculate odds ratio and confidence intervals.

| Environmental variable | Environmental variable | Pr(>\|z\|) |
| --- | --- | --- |
| Location in garden sampled from: |  |  |
| Border (baseline) |  |  |
| + compost bag | 1.64 (0.08-10.32) | 0.65 |
| + compost heap | —<sup>a</sup> | 0.99 |
| + manure bag | —<sup>a</sup> | 0.99 |
| + manure bag + compost heap | —<sup>a</sup> | 0.99 |
| Compost bag | **6.79 (3.25-14.37)** | **<0.01** |
| Compost heap | **4.74 (2.45-9.32)** | **<0.01** |
| Manure bag | —<sup>a</sup> | 0.99 |
| Pot/planter | **2.88 (1.52-5.53)** | **<0.01** |
| + compost bag | **3.05 (1.22-7.26)** | **<0.01** |
| + compost bag + compost heap | —<sup>a</sup> | 0.99 |
| + compost bag + manure bag | —<sup>a</sup> | 0.99 |
| + compost heap | —<sup>a</sup> | 0.99 |

### *Number of tebuconazole-resistant* A. fumigatus *colonies* grown *from soil samples*

The second negative binomial regression was run on the 101 samples that grew tebuconazole-resistant *A. fumigatus*. None of the environmental variables were found to have a significant effect on the outcome.

## Discussion

In this study, 5,174 *A. fumigatus* isolates were cultured from 509 soil samples collected by 249 citizen scientists from their gardens across the UK^44^. Of these soil samples, 327 (64%) grew *A. fumigatus* isolates and 101 (20%) grew isolates that were resistant to tebuconazole at a concentration of 6 mg/L. The percentage of soils that grew *A. fumigatus* in this study was lower than the 78% of soils collected by Sewell *et al.* (2019) from several sites across South West England, including parks, cemeteries, public gardens, flower beds outside hospitals, a lavender farm, a forest and farmland^60^. However, the percentage of soils in this study that grew tebuconazole-resistant *A. fumigatus* isolates was greater than the 6% of soils in Sewell *et al.* (2019) that grew *A. fumigatus* with increased minimum inhibitory concentrations (MICs) to ITZ, VCZ and/or PCZ^60^. Of the 5,174 *A. fumigatus* isolates cultured in this study, 736 (14%) were resistant to tebuconazole, which is greater than the 6% prevalence of triazole-resistant *A. fumigatus* reported by Tsitsopoulou *et al.* (2018) from urban and rural soils in South Wales^61^ and the absence of triazole-resistance detected by van der Torre *et al.* (2020) in isolates cultured from soils adhered to vegetables grown in the UK^62^. This prevalence of 14% is also greater than the 9% in experimental cropland and 12% in commercial wheat fields in the UK reported by Fraaije *et al.* (2020)^57^; however it is less than the 37% prevalence in isolates cultured from flower bulbs bought from a garden centre in Dublin reported by Dunne *et al.* (2017)^63^. In this study, the average concentration of *A. fumigatus* from positive soil samples was 316 CFU/g, which is higher than the 43.5 CFU/g in agricultural soils and 106 CFU/g in urban soils from Greater Manchester reported by Bromley *et al.* (2014)^64^ and considerably higher than the 0-10 CFU/g reported from woodlands, grass verges, experimental cropland and commercial wheat fields across the UK by Fraaije *et al.* (2020)^57^.

Of the 736 *A. fumigatus* isolates that grew on tebuconazole at 6 mg/L, 93 (13%) did not re-grow from short-term storage in the fridge, which left 643 (87%) isolates for sequencing of the *cyp51A* promoter and gene coding regions. Similar to existing UK studies^60,61,64^, the predominant mutation identified in this study was TR_34_/L98H (*n* = 535; 73%). Of these isolates, 22 had amino acid substitutions in cyp51A in addition to L98H. Six isolates had T289A, I364V and G448S amino acid substitutions in addition to TR_34_/L98H; which has been previously detected in Korea in a patient with IA^56^ and in Japan on tulip bulbs imported from The Netherlands^65^. TR68/L98H was detected in one isolate, which was found to be two repeats of the 34-base pair (bp) insert, and in three isolates TR_34_ was detected without any accompanying amino acid substitutions, which was first detected in an environmental isolate collected from Scotland^54^. TR_46_/Y121F/T289A was detected in 16 (2%) isolates and was accompanied by S363P, I364V and G448S in four additional isolates; a combination reported from the Netherlands in 2018^57^. Additional polymorphisms detected in this study included TR_53_, which has been previously reported from flower fields in Colombia^66^ and from a patient with multiple-azole-resistant *A. fumigatus* osteomyelitis in The Netherlands^58^, and TR_92_/Y121F/M172I/T289A/G448S, which has been previously detected in flower bulb waste in The Netherlands^34^ and is two repeats of the 46 bp insert. There were 33 (4%) isolates in this study that did not contain any TRs: five contained I242V, one contained C270R and 27 had no amino acid substitutions in cyp51A. I242V is the only single cyp51A amino acid substitution detected in this study to have been reported in studies summarising *cyp51A* polymorphisms^53,67–69^, which may suggest these polymorphisms occurred *in situ*.

The only environmental variable measured in this study that was found to have a significant effect on whether a sample grew *A. fumigatus*, or on the numbers of *A. fumigatus* grown, was the garden location from which the sample was collected. The greatest concentration of *A. fumigatus* was cultured from a bag of manure at 600 CFU/g, followed by homemade compost heap samples at 505 CFU/g, commercial compost bag samples at 451 CFU/g and pot/planters containing commercial compost at 321 CFU/g. Soil samples that did not contain compost grew fewer *A. fumigatus* isolates: 254 CFU/g from pot/planters and 204 CFU/g from borders. Similar observations were made for tebuconazole-resistant *A. fumigatus*, with concentrations of 128-289 CFU/g recorded for samples containing compost and 98-127 CFU/g for samples without compost. As citizen scientists were only asked to indicate one garden location from which the soil sample was collected, it is possible that the concentrations of *A. fumigatus* spores from borders and pot/planters were inflated by the recent addition of compost that was not indicated on the questionnaire. In the regression models, soils collected from commercial compost bags, homemade compost heaps, pot/planters and pot/planters plus commercial compost had significantly greater odds of growing *A. fumigatus* and tebuconazole-resistant *A. fumigatus* (p < 0.01 for all associations) when compared to soil samples collected from borders. Furthermore, samples collected from commercial compost bags, homemade compost heaps and pot/planter plus commercial compost grew significantly more *A. fumigatus* colonies compared to samples collected from borders. No association was found for garden location sampled from and numbers of tebuconazole-resistant *A. fumigatus*.

Several existing studies have looked for triazole-resistant *A. fumigatus* specifically in compost in the UK and globally, and the findings have been highly variable. Tsitsopoulou *et al.* (2018) collected 11 compost samples from agricultural fields, a horticultural nursery and public areas across South Wales that grew 10 *A. fumigatus* isolates in total; none of which were triazole-resistant^61^. Dunne *et al.* (2017) do not report how many samples they collected from commercial compost bought from a garden centre in Dublin or how many *A. fumigatus* were cultured from these samples; only that one isolate was triazole-resistant^70^. Sewell *et al.* (2019) collected two samples from a compost heap in London that, combined with three samples collected from a flower bed ~500m away, gave a 60% prevalence of triazole-resistant *A. fumigatus*^60^. Pugliese *et al.* (2018) sampled from composting orange peel in Italy and found *A. fumigatus* concentrations of 8.8 × 10^3^ CFU/g at the start of the process rising to 605.7 × 10^3^ CFU/g by the end, yet none of the 30 isolates selected for susceptibility testing were triazole-resistant^71^. Santoro *et al.* (2017) sampled from 11 green and brown composts across Spain, Hungary and Italy and found concentrations of *A. fumigatus* ranging from 100 to 10.6 × 10^3^ CFU/g, yet none of the 30 isolates selected for susceptibility testing were triazole-resistant^72^. Ahangarkani *et al.* (2020) screened isolates cultured from 300 compost samples collected in Iran and detected 57 isolates with elevated MICs to ITZ and VCZ^73^. Zhang *et al.* (2021) collected 114 samples from a plant waste stockpile over 16 months in the Netherlands and detected >10^3^ *A. fumigatus* CFU/g in 74% of samples, with the prevalence of triazole-resistant *A. fumigatus* averaging 50% across all samples^33^. Also in the Netherlands, Schoustra *et al.* (2019) found concentrations of triazole-resistant *A. fumigatus* of 200 CFU/g in household green waste, 1.5-1.8 × 10^3^ CFU/g in compost heaps in residential gardens, up to 2.3 × 10^5^ CFU/g in flower bulb waste and up to 8.4 × 10^4^ CFU/g in organic waste from landscaping^34^.

The key findings of this study are that 64% of soil samples collected from residential gardens in the UK grew *A. fumigatus* and 20% of samples grew tebuconazole-resistant *A. fumigatus*. This means that individuals are very likely to be exposed to both *A. fumigatus* and triazole-resistant *A. fumigatus* spores that are aerosolised from soil when they are undertaking gardening activities^39–43^. Although this study has not undertaken susceptibility testing for the tebuconazole-resistant *A. fumigatus* isolates against medical triazoles, the most commonly detected *cyp51A* polymorphisms TR_34_/L98H and TR_46_/Y121F/T289A are associated with elevated MICs to ITZ, PCZ and VCZ^74^. Furthermore, Hodiamont *et al.* (2009) reported a clinical isolate containing TR_53_ as being resistant to ITR and VCZ, with reduced susceptibility to PCZ^58^. This study also reports that the likelihood of being exposed to *A. fumigatus* and triazole-resistant *A. fumigatus* spores is significantly greater when handling commercial or homemade compost compared to soils in borders or pots/planters. The 14% prevalence of triazole-resistance detected in garden soil samples in this study is higher than most existing studies that have sampled from rural and urban locations in the UK, which is likely being driven by the concentrated application of compost in residential settings. The National Aspergillosis Centre advises that people take care when opening bags of compost and recommends wearing a facemask whilst doing so to avoid dust inhalation. Currently, the only health warning on commercial compost bags is for women to not handle compost without gloves if they are pregnant, presumably to avoid toxoplasmosis infection^75^. The evidence presented here supports the recommendation for users to wear a mask whilst handling compost and the introduction of health warnings on bags of compost with regards to inhaling *A. fumigatus*. Measures could also be taken by compost producers to sterilise the composting before packaging, thereby killing viable *A. fumigatus* spores and eliminating the immediate hazard it poses to the user.

## Acknowledgements

The authors would like to thank all the citizen scientists who collected soil samples for this study. We also thank Dr Pippa Douglas for providing the locations of composters in England with open windrow or outdoor activity, and Dr Jianhua Zhang for sharing the *cyp51A* coding region primers.

## Author Contributions

JMGS, ACS, and MCF conceptualised the study; AA and PSD contributed experimental techniques; JMGS and RC processed samples; JMGS and CU analysed the data; JMGS drafted the original manuscript, which CU, ACS and MCF reviewed and edited.

## References

1. Millner, P. D., Marsh, P. B., Snowden, R. B. & Parr, J. F. Occurrence of Aspergillus fumigatus during composting of sewage sludge. Appl. Environ. Microbiol. 34, 765–772 (1977).

2. Beffa, T. et al. Mycological control and surveillance of biological waste and compost. Med. Mycol. 36, 137–45 (1998).

3. Kwon-Chung, K. J. & Sugui, J. A. Aspergillus fumigatus-What Makes the Species a Ubiquitous Human Fungal Pathogen? PLoS Pathog. 9, 1–4 (2013).

4. Latgé, J. P. Aspergillus fumigatus and Aspergillosis. Clin. Microbiol. Rev. 12, 310–350 (1999).

5. Moss, R. B. Treatment options in severe fungal asthma and allergic bronchopulmonary aspergillosis. Eur. Respir. J. 43, 1487–1500 (2014).

6. Hohl, T. M. & Feldmesser, M. Aspergillus fumigatus: Principles of pathogenesis and host defense. Eukaryot. Cell 6, 1953–1963 (2007).

7. Lin, S., Schranz, J. & Teutsch, S. M. Aspergillosis Case-Fatality Rate?: Systematic Review of the Literature. 60612, (2001).

8. Pegorie, M., Denning, D. W. & Welfare, W. Estimating the burden of invasive and serious fungal disease in the United Kingdom. J. Infect. 74, 60–71 (2017).

9. Bueid, A. et al. Azole antifungal resistance in Aspergillus fumigatus: 2008 and 2009. J. Antimicrob. Chemother. 65, 2116–2118 (2010).

10. van der Linden, J. W. M. et al. Clinical implications of azole resistance in Aspergillus fumigatus, The Netherlands, 2007-2009. Emerg. Infect. Dis. 17, 1846–1854 (2011).

11. Meis, J. F., Chowdhary, A., Rhodes, J. L., Fisher, M. C. & Verweij, P. E. Clinical implications of globally emerging azole resistance in Aspergillus fumigatus. Philos. Trans. B 371, 1–10 (2016).

12. Snelders, E. et al. Triazole fungicides can induce cross-resistance to medical triazoles in Aspergillus fumigatus. PLoS One 7, (2012).

13. Kleinkauf, N. et al. European Centre for Disease Prevention and Control: Risk assessment on the impact of environmental usage of triazoles on the development and spread of resistance to medical triazoles in Aspergillus species. (2013).

14. DEFRA. Government Review of Waste Policy in England 2011. Gov. Waste Policy Rev. 1–80 (2011).

15. HM Government. Food 2030. (2010).

16. Sykes, P., Jones, K. & Wildsmith, J. D. Managing the potential public health risks from bioaerosol liberation at commercial composting sites in the UK: An analysis of the evidence base. Resour. Conserv. Recycl. 52, 410–424 (2007).

17. Deacon, L. et al. Endotoxin emissions from commercial composting activities. Environ. Heal. A Glob. Access Sci. Source 8, 2–5 (2009).

18. Gilbert, E. J., Adrian, K., Karnon, J. D., Swan, J. R. & Crook, B. Preliminary Results of Monitoring the Release of Bioaerosols from Composting Facilities in the UK: Interpretation, Modelling and Appraisal of Mitigation Measures. Biocycle (2002).

19. Sánchez-Monedero, M. A. & Stentiford, E. I. Generation and dispersion of airborne microorganisms from composting facilities. Process Saf. Environ. Prot. 81, 166–170 (2003).

20. Stagg, S., Bowry, A., Kelsey, A. & Crook, B. Bioaerosol emissions from waste composting and the potential for workers’ exposure. Health and Safety Executive (2010).

21. Taha, M. P. M., Drew, G. H., Longhurst, P. J., Smith, R. & Pollard, S. J. T. Bioaerosol releases from compost facilities: Evaluating passive and active source terms at a green waste facility for improved risk assessments. Atmos. Environ. 40, 1159–1169 (2006).

22. Taha, M. P. M., Pollard, S. J. T., Sarkar, U. & Longhurst, P. Estimating fugitive bioaerosol releases from static compost windrows: Feasibility of a portable wind tunnel approach. Waste Manag. 25, 445–450 (2005).

23. Wery, N. Bioaerosols from composting facilities - a review. Front. Cell. Infect. Microbiol. 4, 1–9 (2014).

24. Schantora, A. L. et al. Prevalence of Work-Related Rhino-Conjunctivitis and Respiratory Symptoms Among Domestic Waste Collectors. Environment Exposure to Pollutants (2014).

25. Velasco Garrido, M., Bittner, C., Harth, V. & Preisser, A. M. Health status and health-related quality of life of municipal waste collection workers - A cross-sectional survey. J. Occup. Med. Toxicol. 10, 1–7 (2015).

26. Hoffmeyer, F. et al. Prevalence of and relationship between rhinoconjunctivitis and lower airway diseases in compost workers with current or former exposure to organic dust. Ann. Agric. Environ. Med. 21, 705–711 (2014).

27. Van Kampen, V. et al. Symptoms, spirometry, and serum antibody concentrations among compost workers exposed to organic dust. J. Toxicol. Environ. Heal. - Part A Curr. Issues 75, 492–500 (2012).

28. Hambach, R. et al. Work-related health symptoms among compost facility workers: a cross-sectional study. Arch. Public Heal. 70, 2–6 (2012).

29. Athanasiou, M., Makrynos, G. & Dounias, G. Respiratory health of municipal solid waste workers. Occup. Med. (Chic. Ill). 60, 618–623 (2010).

30. Allmers, H., Huber, H. & Baur, X. Two year follow-up of a garbage collector with allergic bronchopulmonary aspergillosis (ABPA). Am. J. Ind. Med. 37, 438–442 (2000).

31. Poole, C. J. M. & Wong, M. Allergic bronchopulmonary aspergillosis in garden waste (compost) collectors-occupational implications. Occup. Med. (Chic. Ill). 63, 517–519 (2013).

32. Bünger, J., Schappler-Scheele, B., Hilgers, R. & Hallier, E. A 5-year follow-up study on respiratory disorders and lung function in workers exposed to organic dust from composting plants. Int. Arch. Occup. Environ. Health 80, 306–312 (2007).

33. Zhang, J. et al. Dynamics of Aspergillus fumigatus in Azole Fungicide-Containing Plant Waste in the Netherlands (2016-2017). Appl. Environ. Microbiol. 87, 1–12 (2021).

34. Schoustra, S. E. et al. Environmental hotspots for azole resistance selection of Aspergillus fumigatus, the Netherlands. Emerg. Infect. Dis. 25, (2019).

35. Oxford Economics. The Economic Impact of Ornamental Horticulture and Landscaping in the UK - A Report for the Ornamental Horticulture Round Table Group. (2018).

36. Déportes, I., Benoit-Guyod, J. L. & Zmirou, D. Hazard to man and the environment posed by the use of urban waste compost: a review. Sci. Total Environ. 172, 197–222 (1995).

37. Defra. Household Waste Prevention Evidence Review-A report for Defra’s Waste and Resources Evidence Programme. (2009).

38. Brown, J. E., Masood, D., Couser, J. I. & Patterson, R. Hypersensitivity pneumonitis from residential composting: residential composter’s lung. Ann. allergy, asthma Immunol. 74, 45–7 (1995).

39. Cavling Arendrup, M., Ronan O’Driscoll, B., Petersen, E. & Denning, D. W. Acute pulmonary aspergillosis in immunocompetent subjects after exposure to bark chippings. Scand. J. Infect. Dis. 38, 945–949 (2006).

40. Batard, E., Renaudin, K., Morin, O., Desjars, P. & Germaud, P. Fatal acute granulomatous pulmonary aspergillosis in a healthy subject after inhalation of vegetal dust. Eur. J. Clin. Microbiol. Infect. Dis. 22, 357–359 (2003).

41. Jung, N. et al. Gardening can induce pulmonary failure: Aspergillus ARDS in an immunocompetent patient, a case report. BMC Infect. Dis. 14, 1–3 (2014).

42. Russell, K., Broadbridge, C., Murray, S., Waghorn, D. & Mahoney, A. Gardening can seriously damage your health. Lancet 371, 2056 (2008).

43. Zuk, L. A., King, D., Zakhour, H. D. & Delaney, J. C. Locally invasive pulmonary aspergillosis occurring in a gardener: an occupational hazard? Thorax 44, 678–679 (1989).

44. Shelton, J. M. G., Fisher, M. C. & Singer, A. C. Campaign-Based Citizen Science for Environmental Mycology: The Science Solstice and Summer Soil-Stice Projects to Assess Drug Resistance in Air- and Soil-Borne *Aspergillus fumigatus*. Citiz. Sci. Theory Pract. 5, 20 (2020).

45. Boyle, D. G., Boyle, D. B., Olsen, V., Morgan, J. A. T. & Hyatt, A. D. Rapid quantitative detection of chytridiomycosis (Batrachochytrium dendrobatidis) in amphibian samples using real-time Taqman PCR assay. Dis. Aquat. Organ. 60, 141–148 (2004).

46. Met Office. HadUK-Grid datasets. Available at: https://www.metoffice.gov.uk/research/climate/maps-and-data/data/haduk-grid/datasets. (Accessed: 27th January 2021)

47. UKCEH: Land Cover Map 2019. Available at: https://catalogue.ceh.ac.uk/documents/31f4887a-1691-4848-b07c-61cdc468ace7. (Accessed: 27th January 2021)

48. QGIS. Available at: https://www.qgis.org/en/site/. (Accessed: 27th January 2021)

49. Scottish Environment Protection Agency (SEPA): Waste Sites. (2019). Available at: https://www.sepa.org.uk/data-visualisation/waste-sites-and-capacity-tool/. (Accessed: 4th May 2021)

50. Natural Resources Wales: Environmental Permitting Regulations – Waste Sites. (2021). Available at: http://lle.gov.wales/catalogue/item/EnvironmentalPermittingRegulationsWasteSites/?lang=en. (Accessed: 4th May 2021)

51. Northern Ireland Environment Agency: Waste Licenses Register. Available at: https://appsd.daera-ni.gov.uk/wastelicences/. (Accessed: 4th May 2021)

52. Team, R. C. R: A language and environment for statistical computing. R Foundation for Statistical Computing, Vienna, Austria. (2020). Available at: https://www.r-project.org/.

53. Bernal-Martínez, L., Alastruey-Izquierdo, A. & Cuenca-Estrella, M. Diagnostics and susceptibility testing in Aspergillus. Future Microbiol. 11, 315–328 (2016).

54. Rhodes, J. et al. Tracing patterns of evolution and acquisition of drug resistant Aspergillus fumigatus infection from the environment using population genomics. bioRxiv 1–57 (2021).

55. Snelders, E. et al. Emergence of azole resistance in Aspergillus fumigatus and spread of a single resistance mechanism. PLoS Med. 5, 1629–1637 (2008).

56. Cho, S. Y. et al. Epidemiology and antifungal susceptibility profile of aspergillus species: Comparison between environmental and clinical isolates from patients with hematologic malignancies. J. Clin. Microbiol. 57, 1–13 (2019).

57. Fraaije, B., Atkins, S., Hanley, S., Macdonald, A. & Lucas, J. The Multi-Fungicide Resistance Status of Aspergillus fumigatus Populations in Arable Soils and the Wider European Environment. Front. Microbiol. 11, 1–17 (2020).

58. Hodiamont, C. J. et al. Multiple-azole-resistant Aspergillus fumigatus osteomyelitis in a patient with chronic granulomatous disease successfully treated with long-term oral posaconazole and surgery. Med. Mycol. 47, 217–220 (2009).

59. Majima, H., Teppei, A., Watanabe, A. & Kamei, K. D430G mutation of cyp51A in Aspergillus fumigatus causes azole-resistance. in 9th Advances Against Aspergillosis and Mucormycosis 90 (2020).

60. Sewell, T. R. et al. Elevated Prevalence of Azole-Resistant Aspergillus fumigatus in Urban versus Rural Environments in the United Kingdom. Antimicrob. Agents Chemother. 63, 1–8 (2019).

61. Tsitsopoulou, A. et al. Determination of the prevalence of triazole resistance in environmental Aspergillus fumigatus strains isolated in South Wales, UK. Front. Microbiol. 9, 1–8 (2018).

62. van der Torre, M. H. et al. Absence of azole antifungal resistance in Aspergillus fumigatus isolated from root vegetables harvested from UK arable and horticultural soils. J. Fungi 6, 1–10 (2020).

63. Dunne, K., Hagen, F., Pomeroy, N., Meis, J. F. & Rogers, T. R. Intercountry Transfer of Triazole-Resistant Aspergillus fumigatus on Plant Bulbs. Clin. Infect. Dis. 65, 147–149 (2017).

64. Bromley, M. J. et al. Occurrence of azole-resistant species of Aspergillus in the UK environment. J. Glob. Antimicrob. Resist. 2, 276–279 (2014).

65. Hagiwara, D. Isolation of azole-resistant Aspergillus fumigatus from imported plant bulbs in Japan and the effect of fungicide treatment. J. Pestic. Sci. 45, 147–150 (2020).

66. Alvarez-Moreno, C. et al. Azole-resistant Aspergillus fumigatus harboring TR34/L98H, TR46/Y121F/T289A and TR53 mutations related to flower fields in Colombia. Sci. Rep. 7, (2017).

67. Rybak, J. M., Fortwendel, J. R. & Rogers, P. D. Emerging threat of triazole-resistant Aspergillus fumigatus. J. Antimicrob. Chemother. 74, 835–842 (2019).

68. Howard, S. J. et al. Frequency and evolution of azole resistance in Aspergillus fumigatus associated with treatment failure. Emerg. Infect. Dis. 15, 1068–1076 (2009).

69. van der Torre, M. H., Novak-Frazer, L. & Rautemaa-Richardson, R. Detecting azole-antifungal resistance in Aspergillus fumigatus by pyrosequencing. J. Fungi 6, 1–15 (2020).

70. Dunne, K., Hagen, F., Pomeroy, N., Meis, J. F. & Rogers, T. R. Intercountry Transfer of Triazole-Resistant Aspergillus fumigatus on Plant Bulbs. Clin. Infect. Dis. 65, (2017).

71. Pugliese, M., Matić, S., Prethi, S., Gisi, U. & Gullino, M. L. Molecular characterization and sensitivity to demethylation inhibitor fungicides of Aspergillus fumigatus from orange-based compost. PLoS One 13, 1–18 (2018).

72. Santoro, K. et al. Abundance, genetic diversity and sensitivity to demethylation inhibitor fungicides of Aspergillus fumigatus isolates from organic substrates with special emphasis on compost. Pest Manag. Sci. 73, 2481–2494 (2017).

73. Ahangarkani, F. et al. First azole-resistant Aspergillus fumigatus isolates with the environmental TR46/Y121F/T289A mutation in Iran. Mycoses 63, 430–436 (2020).

74. Buil, J. B., Hagen, F., Chowdhary, A., Verweij, P. E. & Meis, J. F. Itraconazole, voriconazole, and posaconazole CLSI MIC distributions for wild-type and azole-resistant Aspergillus fumigatus isolates. J. Fungi 4, 1–9 (2018).

75. Jones, J. L., Krueger, A., Schulkin, J. & Schantz, P. M. Toxoplasmosis prevention and testing in pregnancy, survey of obstetrician-gynaecologists. Zoonoses Public Health 57, 27–33 (2010).

