## Supplemental Table 1 for "Citizen-science surveillance of triazole-resistant *Aspergillus fumigatus* in UK residential garden soils"

| **Supplementary Table 1:** Details of 736 *Aspergillus fumigatus* isolates cultured from soil samples that grew at tebuconazole 6 mg/L; including the sample they were grown from, the county of the sample location, the garden location from which the sample was collected and the *cyp51A* polymorphisms that were detected. | | | | |
| --- | --- | --- | --- | --- |
| **Sample** | **County, north to south** | **Garden location sampled from** | ***cyp51A* polymorphisms** | **Number of**  ***A. fumigatus* isolates** |
| 077-01 | Aberdeenshire | compost heap | TR_34_/L98H | 12 |
|  |  |  | Failed to re-grow | 1 |
| 077-02 | Aberdeenshire | border | TR_34_/L98H | 1 |
| 057-02 | Perth and Kinross | border | TR_34_/L98H | 4 |
|  |  |  | TR_53_ | 1 |
| 003-01 | Clackmannanshire | compost heap | TR_34_/L98H | 2 |
| 173-01 | Stirlingshire | compost heap | TR_34_/L98H | 55 |
| 086-02 | Edinburgh | pot/planter | TR_34_/L98H | 2 |
|  |  |  | Failed to re-grow | 1 |
| 040-02 | Glasgow | border | TR_34_/L98H | 1 |
|  |  |  | WT | 2 |
|  |  |  | Failed to re-grow | 1 |
| 214-01 | South Lanarkshire | pot/planter | TR_34_/L98H | 2 |
| 214-02 | South Lanarkshire | compost bag | TR_34_/L98H | 4 |
| 226-01 | Belfast | border | Failed to re-grow | 2 |
| 226-02 | Belfast | border | Failed to re-grow | 4 |
| 244-02 | Lancashire | pot/planter | TR_34_/L98H | 3 |
| 045-01 | Lancashire | pot/planter | TR_34_/L98H | 3 |
|  |  |  | I242V | 2 |
|  |  |  | WT | 2 |
|  |  |  | Failed to re-grow | 1 |
| 045-02 | Lancashire | pot/planter | TR_34_/L98H | 5 |
|  |  |  | TR_34_ | 1 |
|  |  |  | I242V | 3 |
|  |  |  | C270R | 1 |
|  |  |  | WT | 13 |
|  |  |  | Failed to re-grow | 2 |
| 068-02 | Lancashire | pot/planter + compost bag | TR_34_/L98H | 1 |
| 137-02 | North Yorkshire | border | TR_34_/L98H | 1 |
| 165-01 | Lincolnshire | compost heap | TR_34_/L98H | 1 |
|  |  |  | TR_46_/Y121F/T289A | 3 |
|  |  |  | (TR_46_)2/Y121F/M172I/T289A/G448S | 1 |
|  |  |  | Failed to re-grow | 3 |
| 126-01 | West Yorkshire | compost bag | TR_34_/L98H | 1 |
| 126-02 | West Yorkshire | compost heap | TR_34_/L98H | 7 |
| 048-02 | Greater Manchester | pot/planter | Failed to re-grow | 6 |
| 024-02 | Greater Manchester | pot/planter | TR_34_/L98H | 3 |
| 182-02 | Greater Manchester | pot/planter + compost bag | TR_34_/L98H/D481N | 1 |
|  |  |  | TR_34_/L98H | 21 |
|  |  |  | Failed to re-grow | 6 |
| 007-02 | Merseyside | border | TR_34_/L98H | 2 |
|  |  |  | Failed to re-grow | 1 |
| 062-02 | Greater Manchester | pot/planter | TR_34_/L98H | 2 |
| 015-02 | Greater Manchester | pot/planter | TR34/L98H/K240R | 1 |
|  |  |  | TR34/L98H/A460S | 1 |
|  |  |  | TR34/L98H | 8 |
|  |  |  | Failed to re-grow | 1 |
| 015-03 | Greater Manchester | border | TR_34_/L98H/K240R | 1 |
|  |  |  | TR_34_/L98H | 29 |
| 015-05 | Greater Manchester | border | Failed to re-grow | 1 |
| 015-16 | Greater Manchester | pot/planter | TR_34_/L98H | 4 |
| 198-02 | South Yorkshire | compost heap | TR_34_/L98H | 1 |
| 129-01 | South Yorkshire | compost bag | TR_34_/L98H | 2 |
|  |  |  | Failed to re-grow | 2 |
| 106-01 | Cheshire | pot/planter | TR_34_/L98H | 2 |
| 102-01 | Derbyshire | pot/planter | *Aspergillus fumigatus* - failed to sequence | 1 |
| 023-02 | Cheshire | compost heap | TR_34_/L98H | 17 |
| 167-02 | Cheshire | compost bag | TR_34_/L98H | 2 |
| 074-02 | Lincolnshire | border | TR_34_/L98H | 11 |
|  |  |  | (TR_34_)^2^/L98H | 1 |
|  |  |  | Failed to re-grow | 18 |
| 078-01 | Nottinghamshire | pot/planter | TR_34_/L98H | 4 |
|  |  |  | WT | 5 |
|  |  |  | Failed to re-grow | 5 |
| 216-01 | Lincolnshire | pot/planter | TR_34_/L98H | 1 |
| 231-01 | Nottinghamshire | compost heap | TR_34_/L98H | 1 |
| 160-01 | Norfolk | compost heap | TR_34_/L98H | 26 |
|  |  |  | Failed to re-grow | 4 |
| 160-02 | Norfolk | compost bag | TR_34_/L98H | 9 |
| 255-01 | Shropshire | compost heap | TR_34_/L98H/K372R | 1 |
|  |  |  | TR_34_/L98H | 12 |
|  |  |  | TR_46_/Y121F/T289A | 1 |
|  |  |  | Identity unknown – failed to sequence | 1 |
| 210-01 | Norfolk | pot/planter + compost bag | TR_34_/L98H | 2 |
| 207-01 | West Midlands | compost bag | TR_34_/L98H/D481N | 1 |
|  |  |  | TR_34_/L98H | 5 |
|  |  |  | Failed to re-grow | 6 |
| 018-02 | West Midlands | border | TR_34_/L98H | 2 |
| 101-01 | Worcestershire | border | TR_34_/L98H | 10 |
| 082-02 | Cambridgeshire | compost heap | TR_46_/Y121F/T289A | 1 |
| 032-01 | Cambridgeshire | pot/planter | TR_34_/L98H/P443L | 1 |
|  |  |  | TR_34_/L98H | 2 |
|  |  |  | WT | 2 |
| 054-02 | Essex | border | TR_34_/L98H | 3 |
|  |  |  | WT | 1 |
|  |  |  | Failed to re-grow | 1 |
| 135-01 | Gloucestershire | pot/planter | TR_34_/L98H | 6 |
|  |  |  | Failed to re-grow | 2 |
| 135-02 | Gloucestershire | pot/planter | TR_34_/L98H | 1 |
|  |  |  | Failed to re-grow | 2 |
| 097-01 | Gloucestershire | pot/planter + compost bag | TR_34_/L98H | 6 |
| 149-01 | Hertfordshire | pot/planter + compost bag | TR_34_/L98H | 2 |
|  |  |  | TR_34_ | 1 |
|  |  |  | TR_46_/Y121F/T289A | 1 |
|  |  |  | Failed to re-grow | 5 |
| 149-02 | Hertfordshire | pot/planter | TR_34_/L98H | 1 |
| 055-02 | Oxfordshire | compost heap | TR_34_/L98H/T289A/I364V/G448S | 1 |
|  |  |  | TR_34_/L98H | 28 |
|  |  |  | TR_46_/Y121F/T289A | 1 |
| 038-01 | Oxfordshire | compost bag | TR_34_/L98H/P394R | 1 |
|  |  |  | TR_34_/L98H/K421R | 1 |
|  |  |  | TR_34_/L98H | 28 |
| 150-01 | Pembrokeshire | border | TR34/L98H | 4 |
|  |  |  | Failed to re-grow | 1 |
| 150-02 | Pembrokeshire | compost heap | Failed to re-grow | 1 |
| 158-02 | Buckinghamshire | compost heap | TR34/L98H/T289A/I364V/G448S | 5 |
|  |  |  | TR34/L98H | 7 |
|  |  |  | TR46/Y121F/T289A | 3 |
|  |  |  | TR53 | 1 |
|  |  |  | *Aspergillus fumigatus –* failed to sequence | 1 |
|  |  |  | Failed to re-grow | 2 |
| 146-02 | Buckinghamshire | compost bag | Failed to re-grow | 3 |
| 072-01 | Oxfordshire | compost heap | TR_34_/L98H | 1 |
| 033-02 | Mid Glamorgan | compost bag | TR_34_/L98H | 8 |
| 071-02 | Essex | compost bag | TR_34_/L98H | 2 |
| 009-01 | Oxfordshire | compost heap | TR_34_/L98H/N406M | 1 |
|  |  |  | TR_34_/L98H | 3 |
|  |  |  | WT | 1 |
|  |  |  | Failed to re-grow | 3 |
| 009-02 | Oxfordshire | pot/planter | TR_34_/L98H | 1 |
|  |  |  | Failed to re-grow | 1 |
| 029-01 | Oxfordshire | pot/planter | TR_34_/L98H | 2 |
| 029-02 | Oxfordshire | compost bag | TR_34_/L98H/F404V | 1 |
|  |  |  | TR_34_/L98H | 5 |
| 151-01 | Greater London | pot/planter + compost bag | Failed to re-grow | 3 |
| 229-01 | Berkshire | compost heap | TR_46_/Y121F/T289A | 1 |
| 084-02 | Oxfordshire | pot/planter | TR_34_/L98H | 1 |
|  |  |  | WT | 1 |
| 123-02 | Greater London | pot/planter | *Aspergillus fischeri* | 5 |
| 134-01 | South Gloucestershire | compost bag | TR_34_/L98H | 1 |
| 253-01 | Surrey | border | TR_34_/L98H | 1 |
| 099-01 | Bristol | pot/planter | TR_34_/L98H | 4 |
| 249-01 | Berkshire | compost heap | TR_34_/L98H | 2 |
| 161-01 | Kent | compost heap | TR_34_/L98H | 1 |
| 130-01 | Greater London | border | Failed to re-grow | 1 |
| 010-01 | Greater London | pot/planter | TR_34_/L98H/P443L | 1 |
|  |  |  | TR_34_/L98H | 6 |
|  |  |  | Failed to re-grow | 2 |
| 010-02 | Greater London | compost bag | TR_34_/L98H/F404C/F459S/A460S | 1 |
|  |  |  | TR_34_/L98H/N406D | 1 |
|  |  |  | TR_34_/L98H | 3 |
| 087-01 | Greater London | border | *Aspergillus oerlinghausensis* | 3 |
| 087-02 | Greater London | compost heap | TR_34_ | 1 |
|  |  |  | TR_34_/L98H | 8 |
|  |  |  | TR_46_/Y121F/T289A/S363P/I364V/G448S | 1 |
|  |  |  | TR_46_/Y121F/T289A | 3 |
|  |  |  | (TR_130_)^3^/D430G | 4 |
| 257-02 | Berkshire | compost bag | TR_34_/L98H | 3 |
|  |  |  | TR_46_/Y121F/T289A | 1 |
| 056-02 | Greater London | compost bag | TR_34_/L98H | 30 |
| 195-02 | Wiltshire | border | TR_34_/L98H | 12 |
| 039-01 | Somerset | compost heap | TR_34_/L98H | 3 |
| 127-01 | Greater London | pot/planter + compost bag | TR_34_/L98H | 1 |
| 127-02 | Greater London | compost heap | TR_34_/L98H | 1 |
|  |  |  | TR_46_/Y121F/T289A | 1 |
| 053-01 | Surrey | compost heap | TR_34_/L98H | 3 |
| 012-02 | Surrey | compost bag | TR_34_/L98H | 4 |
| 052-01 | Kent | border | TR_34_/L98H | 1 |
| 052-02 | Kent | pot/planter | *Aspergillus fischeri* | 2 |
| 114-02 | Devon | compost bag | TR_34_/L98H/Q191E | 1 |
|  |  |  | TR_34_/L98H | 3 |
| 136-02 | Somerset | compost bag | TR_34_/L98H | 2 |
|  |  |  | Failed to re-grow | 1 |
| 110-02 | Devon | compost heap | TR_34_/L98H | 3 |
|  |  |  | *Aspergillus fischeri* | 1 |
| 256-01 | Devon | border/compost bag | TR_34_/L98H/R196L | 1 |
|  |  |  | TR_34_/L98H | 7 |
| 238-02 | Hampshire | pot/planter | TR_34_/L98H | 1 |
| 073-01 | Sussex | compost heap | TR_46_/Y121F/M178W/T289A/S363P/I364V/G448S | 1 |
|  |  |  | TR_46_/Y121F/T289A/S363P/I364V/G448S | 3 |
| 091-02 | Sussex | pot/planter + compost bag | TR_34_/L98H | 1 |
| 199-01 | Hampshire | compost heap | TR_34_/L98H | 30 |
| 107-02 | Dorset | compost bag | TR_34_/L98H | 2 |
| 042-01 | Isle of Wight | compost bag | TR_34_/L98H | 3 |
| 176-01 | Cornwall | compost heap | TR_34_/L98H | 13 |
| 124-01 | Devon | pot/planter | TR_34_/L98H | 1 |
